# A toolkit to characterize protein polymerization from cryo-electron tomography data

**DOI:** 10.1101/2025.04.22.650024

**Authors:** Forrest Lee, Victor Manuel Fenton Aguilar, Tripp Lawrence, Philip Mancino, Staas Lin, Kristy Rochon, Lauren Ann Metskas

## Abstract

**Background:** Cryogenic electron tomography (cryo-ET) is a powerful method to study protein structures and macromolecular complexes. These studies can provide structural information at the nanometer scale, allowing for the visualization of ultrastructures and access to subnanometer information through localized, subtomogram averaging (STA). STA alignments can provide analysts with an opportunity to quantify relationships between particles; however, the analytical tools to accomplish this are often lab or system specific. We offer a robust MATLAB script package that can be applied to a wide array of systems.

**Methods:** Many tomographic analyses require extensive segmentation and image classification, which though useful can have poorly characterized error rates and user biases. We present a method for unbiased, numerical classification of head-to-tail polymerization in STA datasets with no outside information or user influence required. We provide this code in a modular MATLAB script package for ease of adaptation to other projects, with analyses including volumes, concentrations, binding, and fibril bundling.

**Results:** We demonstrate the robust analysis possible with this script package using a model system of Rubisco in α-carboxysomes (α-CBs), showcasing the code’s ability to evaluate global data such as volume and overall organization, polymerization data such as twist and bend, and lattice data such as lateral fibril distances and angles.

**Discussion:** Our script package offers structural biologists a toolkit to conduct a robust biophysical analysis on STA data in an unbiased manner. The information generated will provide new insights into protein-protein interactions and the conditions favorable for larger ultrastructures. Particles can also be classified for further STA processing. This script package can be used for scientists studying proteins within isolated compartments or with clearly defined regions of interest.

## Introduction

Cryogenic electron tomography (cryo-ET) provides three-dimensional imaging of cells, tissue slices, and purified samples; subtomogram averaging (STA) extends tomography resolution by aligning and averaging subvolumes within the images to allow protein structure determination at angstrom to nanometer resolution. Although protein structures solved with STA are typically lower resolution than those solved with cryogenic electron microscopy (cryo-EM) single particle averaging, they provide additional contextual information such as cellular localization or interactions with other proteins [1–3].

STA is particularly powerful in evaluating protein ultrastructures, where the arrangement of individual subunits can be determined and the organizational principles derived from the protein positions [4–6]. In recent years several approaches have been developed to increase information extraction from tomograms beyond protein structures, such as shape analyses [7]. Many of these approaches are lab- and project-specific, though recently efforts have been made to provide more generalizable tools to the community [8].

Here we present a script library for analyzing fibrils directly from STA data. Unlike approaches based on image classification, our numerical analysis allows a scientist to control critical parameters and propagate error, enabling rigorous biophysical analysis such as the calculation of binding affinities and nucleation factors. It is also more computationally efficient than image classification. While the pseudocode for this script library was developed to analyze end-to-end polymerization [9], its new modular nature, generalized inputs and new underlying calculations now allow for its use in sorting monomers and dimers, or different layers of a cluster of monomers, enabling a wider range of applications.

## Method

### Overview

Our method relies on accurate knowledge of the positions and orientations of target monomers in a region of interest. As such, a STA dataset with accurate particle picking and resolved α-helices is required to begin this analysis. We have previously published a pipeline for subtomogram averaging that preserves biophysical information while achieving the necessary resolution [10]. Our script package utilizes the Dynamo table format for data input, though these table files can be prepared from the results of any major STA software [11]. The user inputs the table and basic information about the project to the main pipeline file, performs a guided analysis, and then has a range of options for further evaluation (Fig. 1).

**Figure 1.**
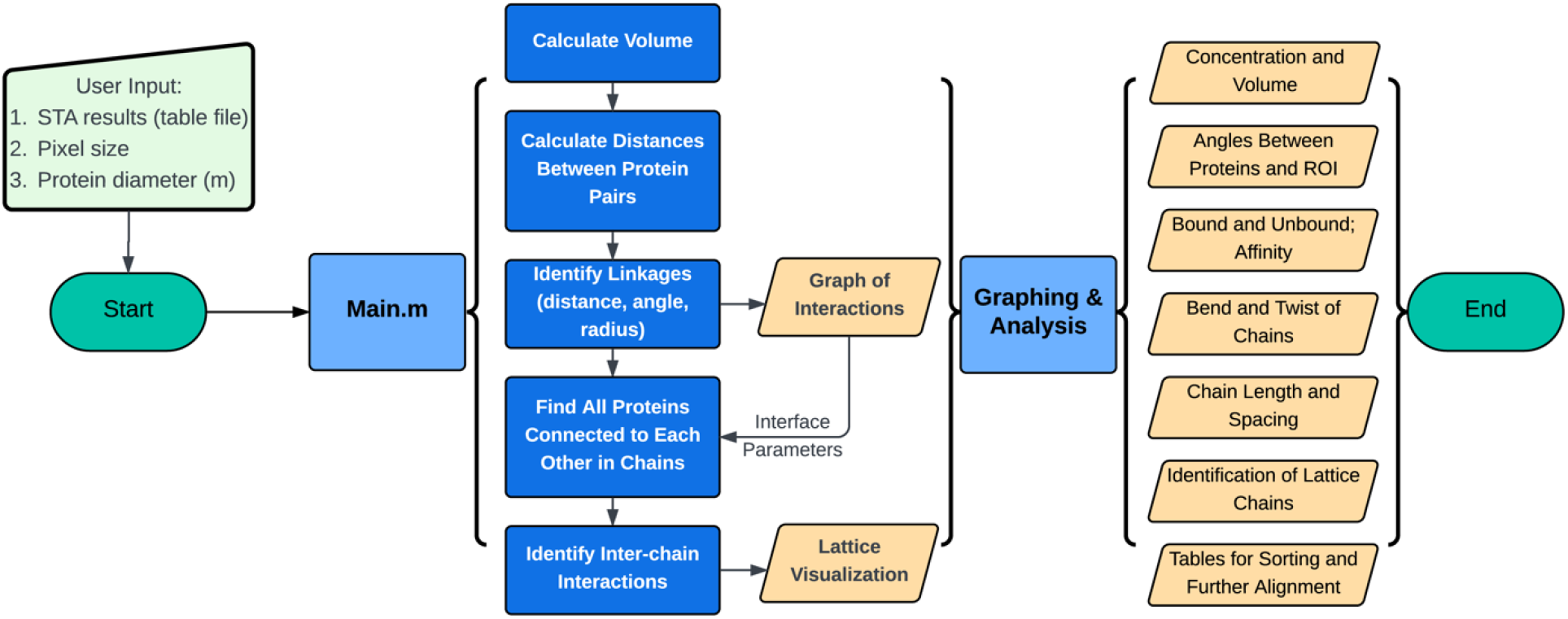
User input and toolkit output workflow. The toolkit is executed through one main MATLAB script (main.m) which calls the core functions. After the main script is executed, a package of graphing and analysis modules are available to evaluate biophysical properties of the particles and the system.

We begin by defining a region of interest (ROI) containing a group of particles (Fig. 2A). This may be determined by the biology, such as an intact virus, microcompartment, or other entity; otherwise, the ROI can be any cluster of particles for which an analysis should be performed. The particles belonging to the ROI are grouped and the volume and concentration of the group are calculated.

**Figure 2.**
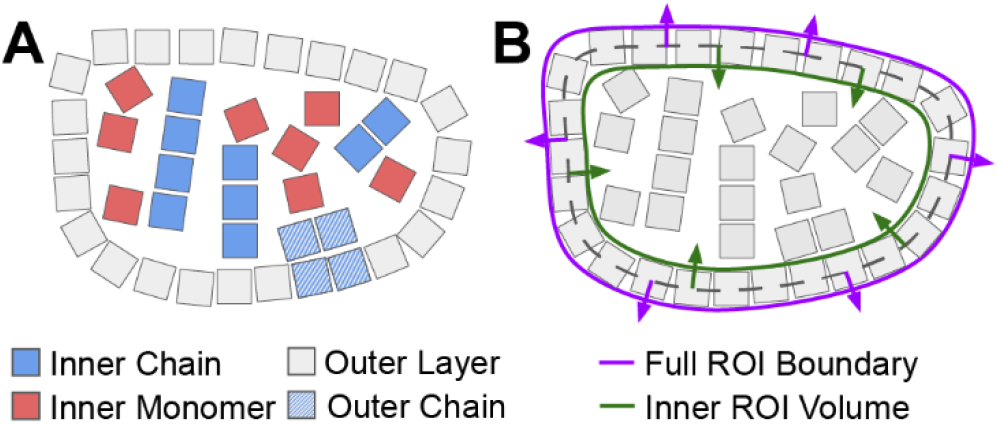
Schematic of an ROI with several possible configurations of the target particle which our script package is designed to identify. **A** Target particles are indicated as boxes. **B** The ROI boundary is determined by a convex hull using the outer particle layer (dashed line). The full ROI boundary is expanded outward to include all particles (purple line); the inner ROI volume is calculated by contracting the boundary inward (green line) to eliminate the outermost layer.

We next build linkages between the particles within an ROI. Particles are linked if they lie within specific distance and angle constraints that suggest polymerization or other targeted interaction types. Particles without a linkage are classified as monomers; transitively bound particles become chains. We then establish the unique order of the linkages within the chain based on their position in the ROI frame of reference. Once chains are established, we look for bundling according to the distance between chains, and group bundles with members in common. Finally, the full ROI is categorized based on the properties of the bundles it contains. Parameters for defining linkages, chains, and bundles are all determined directly from the data, though the analyst may instead choose to set bounds based on other sources of biochemical or structural information.

Other scripts in the package use the calculations and classifications described above to conduct further analyses, including creating binding curves, orientation parameter plots, and chain twist histograms (Fig. 1).

### ROI volume and concentration

We use a built-in MATLAB function to perform a convex hull analysis on the particles in each ROI to define the occupied space, eliminating the need for independently defined boundaries. The ROI volume is calculated by defining a random particle inside the ROI as a center, calculating the volume of a segment formed by the center and three ROI boundary particles, and adding all the segments [9]. Boundaries are often unique in biological systems; therefore, we note position by defining particles participating in the convex hull as “outer” and the rest “inner” to enable the user to include or exclude them for various analyses at their discretion. The concentration is calculated by counting particles per volume.

Because the particle positions define the center of the particle, a simple volume calculation using these positions would underestimate the boundary. Therefore, we calculate the normal vectors of triangles formed by three adjacent particles in the convex hull, and expand the convex hull in the direction defined by the average of the normal vectors of the triangles adjacent to a given particle (Fig. 2B). The expansion has a magnitude of half the particle diameter multiplied by the square root of 3 (half the length of the long diagonal of a cube with side length of one particle diameter). Movement in the opposite direction (a contraction) excludes the outer particles, generating an “inner” volume (Fig. 2B).

### Defining particle-particle linkages

After the ROI, particle positions and orientations are established, we build particle-particle linkages. To identify chains of interacting subunits (chains) we numerically identify which subunits are interacting based on their relative angles and positions. This provides user control over parameterization and quantifiable false negative and false positives rates for linkages, while being more computationally efficient than popular image classification-based approaches due to the far smaller amount of data being analyzed (three Euler angles and three spatial coordinates, rather than per-pixel density information for subtomogram volumes) [10–12].

Initially we place only two restrictions on interacting subunits: particles A and B should be evaluated for interaction if they belong to the same ROI and are within a user-defined radius (typically two particle diameters). These criteria drastically decrease runtime by reducing the number of possible interactions without being so stringent as to create false negative results. We then calculate three parameters for each potential interacting pair: distance (projection), angle, and offset (radius) (Fig. 3A).

**Figure 3.**
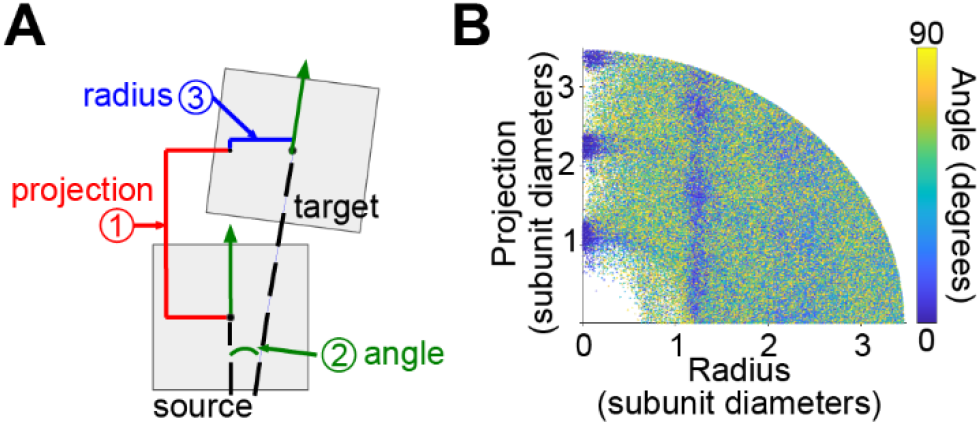
The data are probed to establish subunit interaction parameters. **A** Schematic of a particle pair (boxes), identified based on localization within a preset radius. **B** Every pair of particles has the projection distance (y-axis), radial distance (x-axis), and inter-particle angle (color) defined. Hot spots on this map are used to establish optimal parameters for defining an ordered interaction.

Next, we evaluate the values of these three parameters across the dataset of potential interactions, identifying patterns that reveal how a subunit interaction should be defined (Fig. 3B). Here in the case of polymerization, the strong majority of interactions occur along one axis (x=0), with a step size of approximately one subunit diameter and a small angle between interacting subunits. These high-prevalence values are then turned into parameters by running statistics to find the 2σ thresholds and setting these as the boundaries for defining particles as “bound”. This approach reduces the potential for user bias while allowing the user to track error.

Alternatively, the user can incorporate outside knowledge about their system’s behavior to refine the results. For example, if a structure exists for a chain, the parameters can be extracted from this structure and encoded directly. In such cases, the statistics of Figure 3A could be used to evaluate the number of potential interactions discarded as a result.

The presence of a second set of interactions farther along the Fig. 3B x-axis indicates the presence of lateral interactions for chains (discussed further below).

### Computing chains of linkages

Pairs of interacting subunits with common members are grouped into clusters, and each cluster is used to build one chain (Fig. 4A). The user can specify the minimum number of interacting subunits that defines a chain. All clusters meeting this requirement are then ordered in the original 3D coordinate system of the larger tomogram to give the chain a start and an end. The subunit whose position is furthest from the chain’s center of mass is defined as the start of the chain; all other subunits are then ordered by their Euclidian distance from the starting subunit. Because subunits with D-type symmetry have two identical vector directions due to the mirror plane, we include an optional flipping of vectors in the chain to establish a common direction (Fig. 4A). This allows calculations of the bend and twist of adjacent subunits, without affecting particle orientation in the subtomogram averaging.

**Figure 4.**
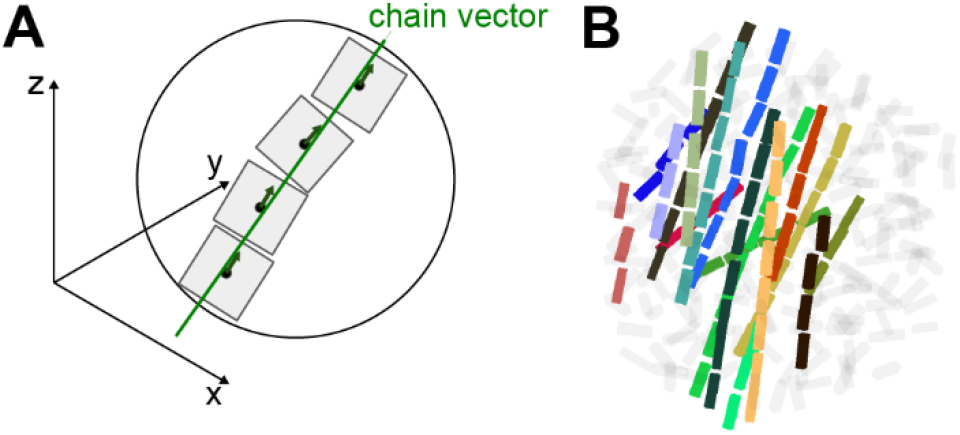
Identifying and organizing longitudinal packing. **A** Continuous chains of three or more particles meeting the interaction parameters are identified and oriented in the 3D space of the original tomogram. **B** A single directional vector is established. **C** Chains can be visualized in the context of the larger ROI including non-chain particles.

We next create a continuous vector to describe the chain and identify its edges. The orientation vectors of each subunit of the chain, now aligned in the same direction, are averaged to obtain a vector that describes the orientation and propagation direction of the chain (Fig. 4A). These chains can be distinguished from monomers and visualized (Fig. 4B) to assess the prevalence and spatial patterning of the chains within the ROI. This visualization can alert an analyst to additional features that may require analysis, such as chain bundling (pictured in Fig. 4B).

### Evaluating lateral packing and fibril bundling

To evaluate lateral packing and bundling, it is necessary to calculate the distance and angle between two chains. We choose not to assume that the chain vector is identical to the chain position, because it is possible that small lateral offsets may have biological significance or that subunits may even shift their offset in response to some inter-chain tether. Therefore, the independence of subunits within a chain is maintained for this calculation. An overlap region is calculated between two chain vectors, and continuous segments are defined within the overlap region as the shortest line connecting two subunit centers (Fig. 5A). The distance between every point on segment for two neighboring chains is then calculated, and the shortest value is reported as the inter-chain distance (derivation in Supplement). The angle between chains is simply the angle between the two chain vectors (see above section).

**Figure 5.**
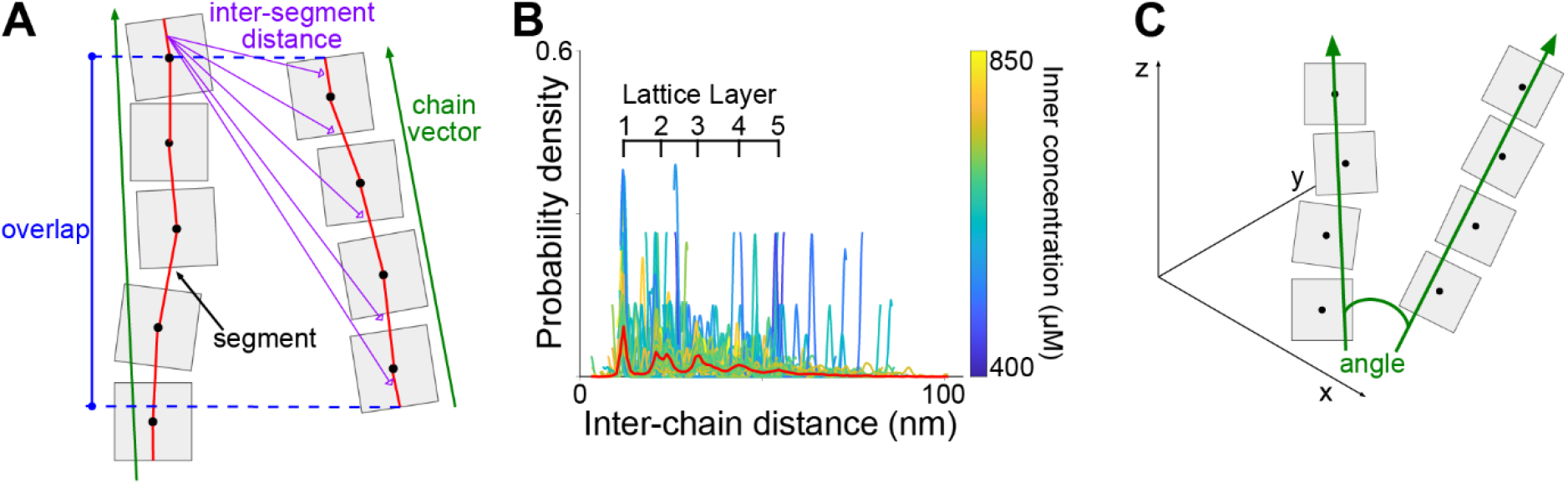
Lateral packing of chains. **A** Schematic of two chains, showing the overlap region (blue) used for calculations, the chain vectors (green), the inter-particle segments (red), and the inter-segment distances (purple). **B** Example probability density plot for a pseudolattice. While there is substantial noise, an average tracing (red line) shows clear peaks at lattice layer positions. For a six-fold arrangement shown here, peaks will begin splitting at layer 2 due to the lattice geometry, and become indistinguishable with increasing distances. **C** Schematic of two chains placed in the reference frame of the tomogram, with the angle calculated from chain vectors (green).

In the case of bundling, it is important to extend inter-chain distances out to multiple lattice layers. This is done easily by adjusting the target distance for the search. A perfect lattice would have discrete distances, while a pseudolattice will show increased propensities for specific distances over a baseline of noise (Fig. 5B).

From this point in the code, general approaches diverge into application-specific analyses (Fig. 1). We provide a series of different, independent options, including volume and concentration estimates, angle of a particle relative to the exterior edge, binding according to concentration, and more. All data are stored in MATLAB objects and the code is fully modular, making it simple for a user to add their own project-specific analyses to the script package.

## Results

We applied this new MATLAB script package to a published, high-accuracy dataset of Rubisco particles polymerizing within α-carboxysomes (α-CBs) [9]. In comparison to the methods used in previous analyses, our core analyses identify subtle changes to calculated Rubisco bend and twist (Fig. 6A-B, Fig. 6D) thanks to improved linkage definitions and inner-layer concentration estimates [9, 13]. The large standard deviation of the fibril twist and non-zero bending angle suggest that the four symmetric binding sites around the Rubisco-Rubisco interface are not all occupied at any given time, allowing some movement between the complexes even as they remain bound at one site. This seems to be a feature of the lattice that allows the symmetry break between the six-fold symmetry of the Rubisco pseudolattice and the four-fold symmetry of the subunits [9].

**Figure 6.**
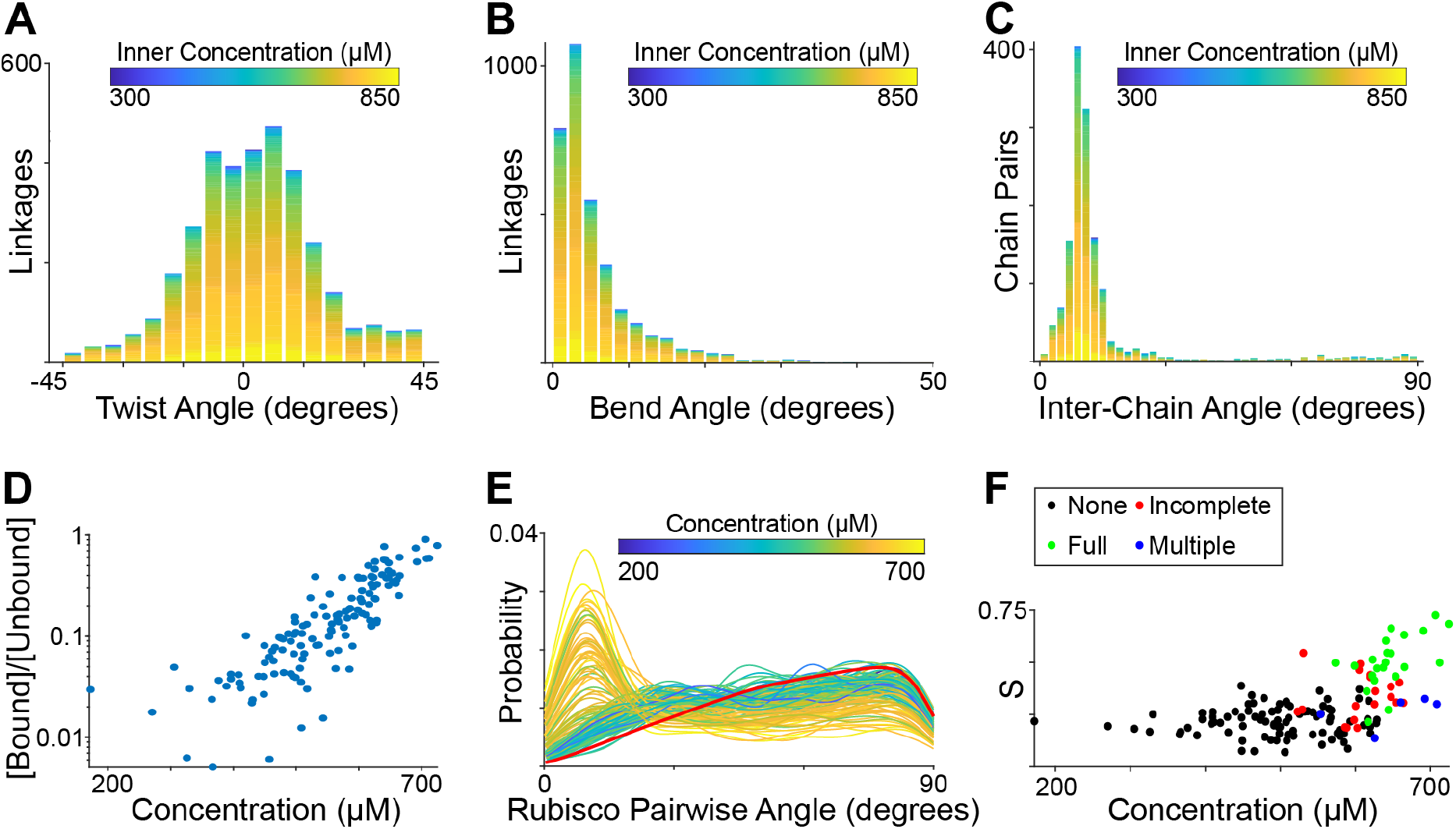
Rubisco inside carboxysomes. Top panels: Rubisco fibrils in the carboxysome interior. **A** Rubisco-Rubisco twist angle and **B** Rubisco-Rubisco bend angle suggest partial occupancy of the four symmetrical binding sites. **C** Fibrils are at a defined angle relative to each other, though this can be broken especially in incomplete or non-canonical pseudolattices. Bottom panels: holistic carboxysome analyses. **D** Rubisco binding curves show a strong concentration dependence for Rubisco polymerization. **E** Nearest-neighbor Rubisco interactions across the interior are dominated by the inter-chain angle, providing information for a second-order tensor calculation. Red line: random behavior. **F** The scalar of the second order tensor showcases the alignment of all Rubisco complexes within one carboxysome and can be used to define different lattice types.

Our added applications include new evaluations of lateral interactions using the continuous chain vectors (Fig. 6C); these applications were not possible with particle-based approaches because the chains might not be in similar registrations (creating severe inconsistencies in inter-chain distances and angles). This also establishes for the first time that the local Rubisco-Rubisco pairwise angles are dominated by the inter-chain angle due to fibril bundling (Fig. 6E). We include a calculation of second-order tensor to evaluate the presence of a lattice, but subdivided into different forms of fibril bundling to identify the carboxysomes where the scalar is artificially reduced due to the presence of multiple lattice seeds pointing in different directions (Fig. 6F). This better clarifies the strong relationship between lattice morphology and concentration compared to previous approaches [9, 13].

For this application, we used initial binding parameters of 0.7 to 1.3 subunit diameters projection distance, a radius of 0.5 subunit diameters, and a maximum angle of 25°, based on an unbiased overview of all near-neighbor interactions (Fig. 3B).

## Discussion

Cryo-EM data have long been used to support binding affinity studies, typically in combination with modeling or free energy calculation. However, attempts to extract rigorous biophysical parameters from cryo-EM and cryo-ET datasets have often failed to control for error in the particle identification and classification steps, making multi-component downstream analyses difficult [10]. Here, we use numerical classification to establish user control over distinctions between “bound” and “unbound” particles, deriving parameters directly from the dataset and giving the user access to error information. The numerical classification is also far more computationally efficient, running over minutes on a small number of CPUs thanks to incorporation of spatial hashing.

Our approach to identifying particle-particle interactions preserves both the particle position and orientation relative to a reference, but also maintains the ultrastructure within the reference space of the tomogram (Fig. 4). This facilitates complex analyses such as sub-classification of lattice types, while preserving the user’s ability to mark particle classes for further STA without having touched the original data. We include several tools in the script library to write Dynamo tables from user-defined parameters, so a user can proceed to run additional STA on subclasses of monomers, chains or specific lattice morphologies. In cases where a small conformation change occurs upon binding, this capability may increase output resolutions by performing a greater-accuracy classification than an image-based approach.

In our tomography test dataset, the target classes (monomer and fibril) are clearly identifiable in the data. If this were not the case, an initial image classification could be performed using existing cryo-EM software or segmentation workflows [11, 14, 15], and this output numerically probed (Fig. 3B) to define parameter sets for numerical classification. We have also provided options to handle multiple symmetry types, including D-type symmetry (requiring vector flips or removal of heads/tails, see Fig. 4).

In conclusion, we have created a modular MATLAB script library with generalized code for evaluation of STA datasets, with particular attention to binding and polymerization. All information is extracted solely from the STA data, reducing user bias and enabling advanced biophysical analyses for which we provide several example functions. We hope that this script package will be readily adaptable to a variety of STA targets as particle picking accuracy improves across the field and more high-accuracy datasets become available.

## Acknowledgements

We thank David Savage and Luke Oltrogge for insightful discussions on carboxysome structure and Rubisco behaviors.

## Authors’ contributions

LAM conceived and supervised the project. FL, LAM and KR wrote the manuscript. FL and VFA wrote the code. FL finalized the toolkit and prepared the software release. SL finalized documentation and GitHub deposition. RL and PM contributed first-generation code.

## Data availability

The source code for this toolkit is available under a non-commercial, share-and-share-alike MIT license at https://github.com/LAMetskas/2025_polymerizationAnalysis/. The original dataset used for Results demonstration is available in EMPIAR, accession number 11125, and EMD-27654.

## References

1. Fäßler F, Javoor MG, Datler J, Döring H, Hofer FW, Dimchev G, et al. ArpC5 isoforms regulate Arp2/3 complex–dependent protrusion through differential Ena/VASP positioning. Science Advances. 2023;9:eadd6495.

2. Böck D, Medeiros JM, Tsao H-F, Penz T, Weiss GL, Aistleitner K, et al. In situ architecture, function, and evolution of a contractile injection system. Science. 2017;357:713–7.

3. Chmielewski D, Schmid MF, Simmons G, Jin J, Chiu W. Chikungunya virus assembly and budding visualized in situ using cryogenic electron tomography. Nat Microbiol. 2022;7:1270–9.

4. Mattei S, Glass B, Hagen WJH, Kräusslich H-G, Briggs JAG. The structure and flexibility of conical HIV-1 capsids determined within intact virions. Science. 2016;354:1434–7.

5. Qu K, Ke Z, Zila V, Anders-Össwein M, Glass B, Mücksch F, et al. Maturation of the matrix and viral membrane of HIV-1. Science. 2021;373:700–4.

6. von Kügelgen A, Tang H, Hardy GG, Kureisaite-Ciziene D, Brun YV, Stansfeld PJ, et al. In Situ Structure of an Intact Lipopolysaccharide-Bound Bacterial Surface Layer. Cell. 2020;180:348–358.e15.

7. Barad BA, Medina M, Fuentes D, Wiseman RL, Grotjahn DA. Quantifying organellar ultrastructure in cryo-electron tomography using a surface morphometrics pipeline. Journal of Cell Biology. 2023;222:e202204093.

8. Jiasuiliu, maschrei314, makubans, turonova, yuningw268, Gioia P, et al. turonova/cryoCAT: New features added. 2025.

9. Metskas LA, Ortega D, Oltrogge LM, Blikstad C, Lovejoy DR, Laughlin TG, et al. Rubisco forms a lattice inside alpha-carboxysomes. Nat Commun. 2022;13:4863.

10. Metskas LA, Wilfong R, Jensen GJ. Subtomogram averaging for biophysical analysis and supramolecular context. J Struct Biol X. 2022;6:100076.

11. Castaño-Díez D, Kudryashev M, Arheit M, Stahlberg H. Dynamo: a flexible, user-friendly development tool for subtomogram averaging of cryo-EM data in high-performance computing environments. J Struct Biol. 2012;178:139–51.

12. Burt A, Toader B, Warshamanage R, von Kügelgen A, Pyle E, Zivanov J, et al. An image processing pipeline for electron cryo-tomography in RELION-5. FEBS Open Bio. 2024;14:1788– 804.

13. Ni T, Sun Y, Burn W, Al-Hazeem MMJ, Zhu Y, Yu X, et al. Structure and assembly of cargo Rubisco in two native α-carboxysomes. Nat Commun. 2022;13:4299.

14. Scheres SHW. RELION: Implementation of a Bayesian approach to cryo-EM structure determination. Journal of Structural Biology. 2012;180:519–30.

15. Heebner JE, Purnell C, Hylton RK, Marsh M, Grillo MA, Swulius MT. Deep Learning-Based Segmentation of Cryo-Electron Tomograms. JoVE (Journal of Visualized Experiments). 2022;:e64435.

